# LCR-modules: a collection of workflows for cancer genome analysis

**DOI:** 10.1101/2025.09.19.676923

**Authors:** Kostiantyn Dreval, Laura K. Hilton, Bruno M. Grande, Krysta M. Coyle, Manuela Cruz, Sierra Gillis, Prasath Pararajalingam, Christopher K. Rushton, Haya Shaalan, Nicole Thomas, Helena Winata, Jasper Wong, Jacky Yiu, Christian Steidl, David W. Scott, Ryan D. Morin

**Author notes:** **Corresponding Author Dr. Ryan D. Morin**, Department of Molecular Biology and Biochemistry, Simon Fraser University, 8888 University Dr., Burnaby, BC, Canada, V5A 1S6.

## Abstract

The surge of genomic data from advanced sequencing technologies is outpacing current analytical pipelines. We introduce LCR-modules, an open-source suite of bioinformatics tools designed for flexible and automated cancer genome data analysis. LCR-modules enables reproducible analysis of diverse cancer genomics data at scale. The suite comprises 49 Snakemake-based workflows organized into three levels, facilitating tasks from low-level quality control to complex cohort-level analyses. LCR-modules supports various sequencing types and integrates pipelines such as mutation calling, expression quantification, and cohort-level aggregation, ensuring flexibility and reproducibility. LCR-modules represents a significant advancement in genomic data analysis, reducing barriers in reproducibility and scalability and has already been applied to a combination of exomes and genomes from over 10,800 samples.

## Introduction

High-throughput sequencing has been transformational to genetic research and the study of cancer (Liu *et al*., 2019; Satam *et al*., 2023; Mustafa, 2024). Large consortia such as The Cancer Genome Atlas(Weinstein *et al*., 2013) and ICGC(Hübschmann *et al*., 2021) and collective work from thousands of teams globally have generated vast amounts of sequencing data from cancer. Generating high-quality simple somatic mutation and copy number alteration calls involves a complex collection of algorithms (Ulrich, 2017; Cremin *et al*., 2022). Bioinformatics workflows commonly rely on a variety of programming languages, which are cumbersome to maintain and difficult to scale as analyses grow in scope and complexity (Dash *et al*., 2019; Wratten *et al*., 2021). Hence, despite the opportunity afforded by a wealth of cancer genomic data to reveal novel molecular insights, the ability to rapidly process data in a reproducible manner serves as a bottleneck. For research to keep pace with the influx of data, pipelines to automate routine analytical steps are essential. Ideally, analytical pipelines balance versatility, tuneability, and ease of installation and maintenance while minimizing risks of errors, thereby enhancing reproducibility. Modularization and version control facilitate the consistent application of analytical best practices and allow deployment of new features and iprovements across projects that share analyses (Davis-Turak *et al*., 2017; Houssein *et al*., 2023). Reliance on modular components helps promote consistency and reproducibility while reducing the amount of lead time in establishing a new analytical framework. To support the routine analyses of short-read exome, panel-based, transcriptome, whole genome sequencing (WGS) and long-read sequencing, we developed a suite of tools collectively named LCR-modules. This toolkit has been used by our team to discover drivers and molecular features in cancers, specifically focusing on lymphoid neoplasms, and perform quantitative comparisons between disease entities (Bal *et al*., 2022; Dreval *et al*., 2022, 2023; Thomas *et al*., 2023, 2025; Hilton *et al*., 2023; Sahasrabuddhe *et al*., 2025). LCR-modules include open-source and custom bioinformatics tools utilizing the Snakemake (Mölder *et al*., 2021) workflow management system, and include modules for data quality control, discovery and annotation of common mutation types, analysis of B-cell receptor repertoires, and discovery of loci affected by aberrant somatic hypermutation. Individual modules are configured to create an automated, scalable, and reproducible workflow that runs each step as dictated by the availability of new data. LCR-modules is actively maintained and openly available on GitHub.

## Results

### LCR-modules overview

LCR-modules is a collection of tools that form the basis of workflows to perform scalable and reproducible genomic analyses. The individual modules can be logically divided into 3 main categories representing processing of raw, sample-level or cohort-level data (**Figure 1A**). The outputs of the modules of each previous level flow, individually or in combination, to modules at the subsequent level. An important feature of the modules at all levels includes a specifically designed set of rules to handle required reference files automatically with minimal effort from the user. These reference rules handle the download of fasta files for majority of the most common genome builds and bed files for commonly used exome panels. While a wide range of genome builds is available to the user out-of-the box, the reference files subworkflow also provides flexibility to add custom reference files or capture-based panels.

**Figure 1.**
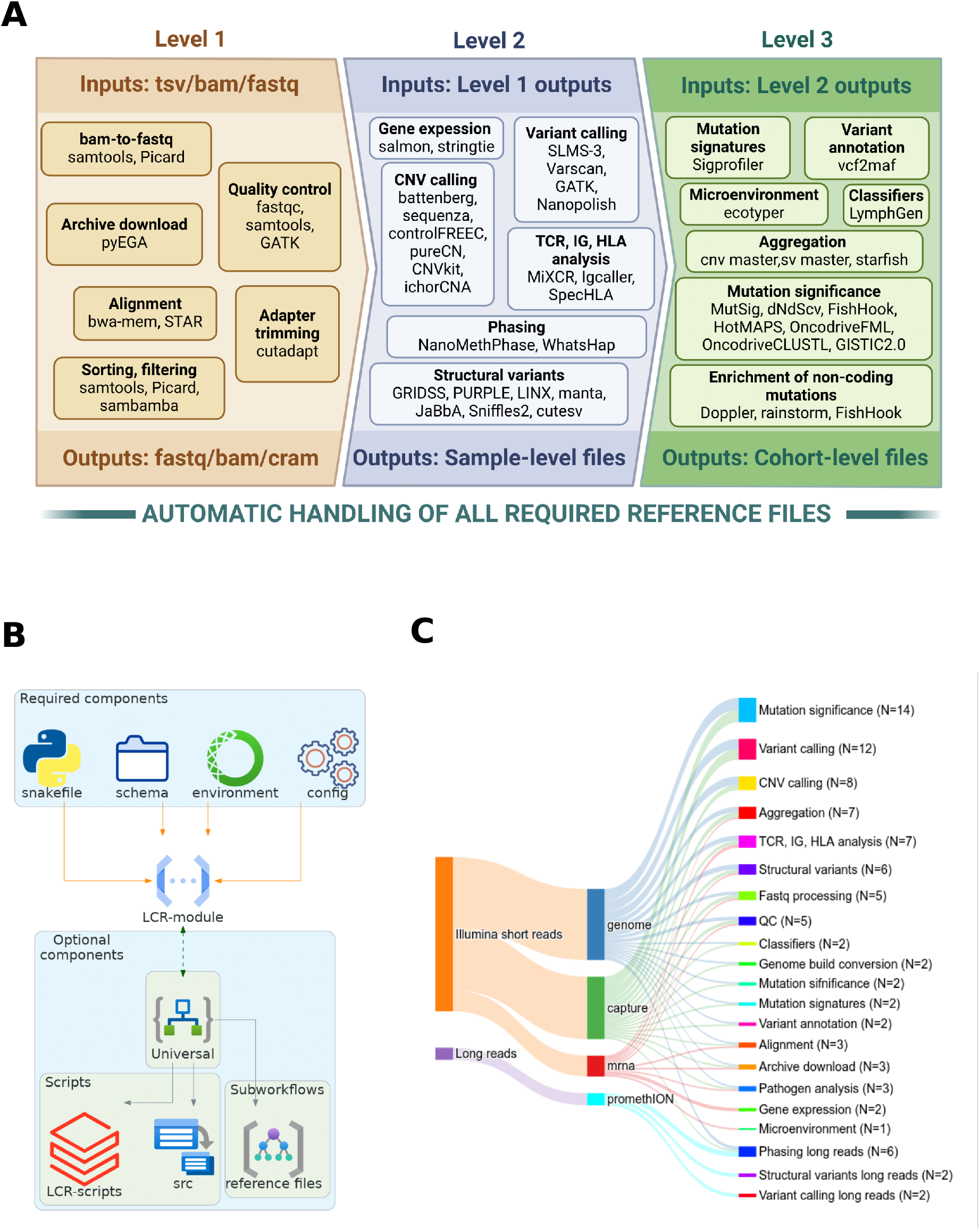
LCR-modules overview. **A**. Schematic representation of modules as organized into different levels according to the complexity of performed analyses. Only representative individual modules are shown. The full list of modules is available at https://github.com/LCR-BCCRC/lcr-modules/tree/master/modules. **B**. The genral structure of main components of individual module. **C**. Sankey plot showing relationship between supported sequencing type and generalized function of each module. The number in parentheses for the left strata illustrates the number of supported modules. The number for the right strata illustrates the combined number of supported sequencing types and individual modules.

Level 1 modules achieve low-level tasks such as adapter trimming, quality control, and alignment of sequencing files and obtaining data from repositories such as the European Genome-phenome Archive (EGA). These modules also perform gene expression analyses, including alignment using (Dobin *et al*., 2013) and calculating mRNA abundance using salmon (Patro *et al*., 2017). Level 2 modules perform routine tasks for cancer genome analysis such as detecting and annotating simple somatic mutations, copy-number alterations and structural variations. Next, the level 3 modules perform analyses that rely on cohort-level aggregation. The cohorts and data sets can be flexibly defined based on different clinical characteristics through a set of configuration files. The modules at this level operate on the outputs of level 2 modules and perform tasks such as aggregation of individual files into cohort-level merges, analyses of mutation signatures, identification of significantly mutated genes, or classification into genetic subgroups. Each module has required and optional components (**Figure 1B**). Required components include: the Snakefile (rules), schema (definition of wildcards), conda environment (dependencies), and module config. The module config defines the input/output paths, tool parameters, computational resource requirements, supported sequencing types (seq_type), and rules for handling data with or without a matched normal. Optional components include a reference subworkflow (automated reference assets with user overrides) and shared custom scripts (provided in-module or via the companion LCR-scripts repository). This structure standardizes behavior, recycles common resources, and minimizes code duplication. The current collection includes 49 workflows designed to support both short-read Illumina and long-read PromethION sequencing data. The scope of supported sequencing data type, or seq_type throughout the LCR-modules suite, currently includes genome, capture, mrna, and long-read sequencing data (Oxford Nanopore). Most of the modules support more than one seq_type, often with automatic configurations tailored to the data (**Figure 1C**). The WGS (genome seq_type) has the highest module support, and the somatic variant calling and identification of significantly mutated genes are the analysis types with the highest representation of the developed modules.

## Assembling modules into pipelines

Any individual modules with compatible inputs or outputs can be combined to facilitate multi-step genomic analysis workflow. Built using Snakemake (Mölder *et al*., 2021), modules can be directly imported into a project Snakefile, exposing their individual rules to the master (project) workflow (**Figure 2A**). To establish a project workflow, each module with its associated config constitutes an individual step, and the individual jobs that will be executed are dictated by the contents of the sample table, the presence of compatible input files, and the target rule definitions. The default configuration values of each module can be replaced with the values specified in the project-level config. Such architecture allows the entire project to be concise, easy to maintain, scalable, and reproducible. The LCR-modules ships with the included demo, where individual modules are organized into a representative workflow based on their sequencing type. The demo project not only supports maintanence of functionality testing for the entire codebase, but also illustrates the concept of the project for users. In addition, the demo contains custom wrapper scripts to launch the workflows as a dry run, for local execution, or for cluster execution using schedulers and workload managers such as Sun Grid Engine and Slurm (Yoo *et al*., 2003). Many of our modules are built atop multiple other modules. One such example is our ensemble tool for detecting simple somatic mutations (SLMS-3). The SLMS-3 workflow independently runs mutect2 (Benjamin *et al*., 2019), strelka2 (Kim *et al*., 2018), SAGE (https://github.com/hartwigmedical/hmftools/tree/master/sage), LoFreq (Wilm *et al*., 2012), and manta (Chen *et al*., 2016) variant callers (**Figure 2B**). Starfish (https://github.com/dancooke/starfish) is used to intersect the resulting variant calls from each individual tool such that only variants supported by 3 or more variant callers are retained. Custom scripts, customized versions of individual callers, and all other required components of SLMS-3 are directly supplied with LCR-modules. This module can further filter variants based on their GnomAD population frequency (Chen *et al*., 2024), minimal sequencing depth (10X), allele frequency (0.1), and other criteria.

**Figure 2.**
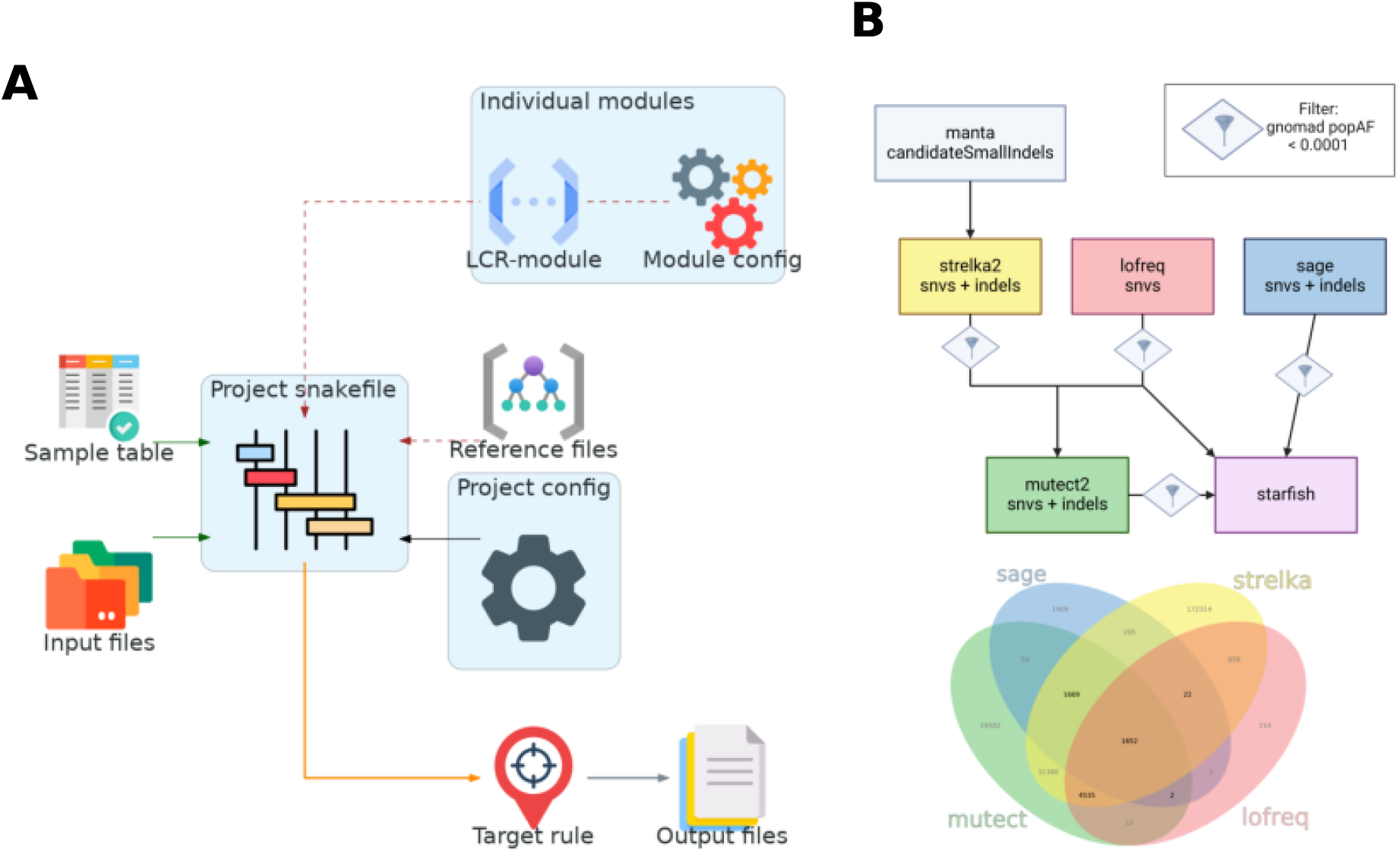
Organizing LCR-modules into a project. **A**. Schematic representation of main components in a project organized with LCR-modules. **B**. Example project for simple somatic varinant calling workflow organized using individual LCR-modules. The venn diagram represents the variants called by each individual tool from a patient with diffuse large B-cell lymphoma (Hilton *et al*., 2023). The numbers on the venn diagram highlighted in bold represent the final set of somatic variants.

We benchmarked the accuracy of SLMS-3 on 44 Burkitt lymphoma biopsies with matched fresh-frozen (FF) and formalin-fixed and paraffin-embedded (FFPE) tissue (Grande *et al*., 2019; Dodani *et al*., 2022; Thomas *et al*., 2023). Somatic variants from the FF samples were recovered in FFPE samples with > 90% sensitivity compared to the variants called in FF samples. When SLMS-3 is deployed on a server with 1500 Gb RAM and 144 Intel(R) Xeon(R) CPU E7-8867 v4 @ 2.40GHz CPUs, it takes 4 h 8 min to complete on a representative paired WGS sample with 40X depth of sequencing, and 1 h 15 min on a representative paired whole exome sample with 53X coverage. Due to the compatibility of LCR-modules with parallel processing on clusters, running SLMS-3 on a real-world data of 100 paired and unpaired FF and FFPE whole exomes from patients with follicular lymphoma (Kalmbach *et al*., 2023) with 135-320X (min-max) depth of coverage takes a total of 3 h 29 min. These execution times highlight fast turnaround time for the analysis of sequencing data and support scalability of the data analysis.

Reproducibility in cancer genomics depends on clear, inspectable pipelines, yet pipeline management is challenged by large datasets, variable genome references and assays, heterogeneous tools, and scaling needs. LCR-modules addresses this with modular Snakemake workflows, pinned conda environments, and a reference subworkflow that automates genome assets. Declarative configs (not ad hoc scripts) control inputs, resources, reference and seq_type handling, reducing developer effort while preserving transparency. Combined with reusable utilities and an included demo for end-to-end testing, this design promotes consistent results across cohorts and infrastructure, and— as shown with SLMS-3—delivers accuracy on FFPE samples while scaling to large studies. To date, LCR-modules has powered analyses in numerous publications (Bal *et al*., 2022; Dreval *et al*., 2022, 2023; Thomas *et al*., 2023, 2025; Hilton *et al*., 2023; Sahasrabuddhe *et al*., 2025) and has processed more than 2060 whole genomes, 4500 exomes, and 4130 RNA-seq datasets — highlighting both its reliability and broad applicability.

## Acknowledgements

This work was supported by a Terry Fox New Investigator Award (No. 1043) and by an operating grant from the Canadian Institutes for Health Research and a New Investigator Award from the Canadian Institutes for Health Research (R.D.M.). R.D.M. is a Michael Smith Foundation for Health Research Scholar, and D.W.S. is a Michael Smith Foundation for Health Research Health Professional-Investigator.

## Data Availability

No new data were generated in support of this research. The source code for the LCR-modules is openly available at https://github.com/LCR-BCCRC/lcr-modules

## References

Bal, E. et al. (2022) Super-enhancer hypermutation alters oncogene expression in b cell lymphoma. Nature, 607, 808–815.

Benjamin, D. et al. (2019) Calling somatic SNVs and indels with Mutect2. BioRxiv, 861054.

Chen, S. et al. (2024) A genomic mutational constraint map using variation in 76,156 human genomes. Nature, 625, 92–100.

Chen, X. et al. (2016) Manta: Rapid detection of structural variants and indels for germline and cancer sequencing applications. Bioinformatics, 32, 1220–1222.

Cremin, C.J. et al. (2022) Big data: Historic advances and emerging trends in biomedical research. Current Research in Biotechnology, 4, 138–151.

Dash, S. et al. (2019) Big data in healthcare: Management, analysis and future prospects. J big data 6: 54.

Davis-Turak, J. et al. (2017) Genomics pipelines and data integration: Challenges and opportunities in the research setting. Expert review of molecular diagnostics, 17, 225– 237.

Dobin, A. et al. (2013) STAR: Ultrafast universal RNA-seq aligner. Bioinformatics, 29, 15–21.

Dodani, D.D. et al. (2022) Combinatorial and machine learning approaches for improved somatic variant calling from formalin-fixed paraffin-embedded genome sequence data. Frontiers in Genetics, 13, 834764.

Dreval, K. et al. (2023) Genetic subdivisions of follicular lymphoma defined by distinct coding and noncoding mutation patterns. Blood, 142, 561–573.

Dreval, K. et al. (2022) Minimal information for reporting a genomics experiment. Blood, The Journal of the American Society of Hematology, 140, 2549–2555.

Grande, B.M. et al. (2019) Genome-wide discovery of somatic coding and noncoding mutations in pediatric endemic and sporadic burkitt lymphoma. Blood, The Journal of the American Society of Hematology, 133, 1313–1324.

Hilton, L.K. et al. (2023) Relapse timing is associated with distinct evolutionary dynamics in diffuse large b-cell lymphoma. Journal of Clinical Oncology, 41, 4164–4177.

Houssein, E.H. et al. (2023) Soft computing techniques for biomedical data analysis: Open issues and challenges. Artificial Intelligence Review, 56, 2599–2649.

Hübschmann, D. et al. (2021) Mutational mechanisms shaping the coding and noncoding genome of germinal center derived b-cell lymphomas. Leukemia, 35, 2002– 2016.

Kalmbach, S. et al. (2023) Novel insights into the pathogenesis of follicular lymphoma by molecular profiling of localized and systemic disease forms. Leukemia, 37, 2058–2065.

Kim, S. et al. (2018) Strelka2: Fast and accurate calling of germline and somatic variants. Nature methods, 15, 591–594.

Liu, Z. et al. (2019) Toward clinical implementation of next-generation sequencing-based genetic testing in rare diseases: Where are we? Trends in Genetics, 35, 852–867.

Mölder, F. et al. (2021) Sustainable data analysis with snakemake. F1000Research, 10.

Mustafa, A.S. (2024) Whole genome sequencing: Applications in clinical bacteriology. Medical Principles and Practice, 33, 185–197.

Patro, R. et al. (2017) Salmon provides fast and bias-aware quantification of transcript expression. Nature methods, 14, 417–419.

Sahasrabuddhe, A.A. et al. (2025) The FBXO45–GEF-H1 axis controls germinal center formation and b-cell lymphomagenesis. Cancer Discovery, 15, 838–861.

Satam, H. et al. (2023) Next-generation sequencing technology: Current trends and advancements. Biology, 12, 997.

Thomas, N. et al. (2025) DNA methylation epitypes of burkitt lymphoma with distinct molecular and clinical features. Blood cancer discovery, 6, 325–342.

Thomas, N. et al. (2023) Genetic subgroups inform on pathobiology in adult and pediatric burkitt lymphoma. Blood, 141, 904–916.

Ulrich, T. (2017) Harnessing the flood: Scaling up data science in the big genomics era. Broad Institute.

Weinstein, J.N. et al. (2013) The cancer genome atlas pan-cancer analysis project. Nature genetics, 45, 1113–1120.

Wilm, A. et al. (2012) LoFreq: A sequence-quality aware, ultra-sensitive variant caller for uncovering cell-population heterogeneity from high-throughput sequencing datasets. Nucleic acids research, 40, 11189–11201.

Wratten, L. et al. (2021) Reproducible, scalable, and shareable analysis pipelines with bioinformatics workflow managers. Nature methods, 18, 1161–1168.

Yoo, A.B. et al. (2003) Slurm: Simple linux utility for resource management. In, Workshop on job scheduling strategies for parallel processing. Springer, pp. 44–60.

